# OBSTRUCTIVE APNEAS IN A MOUSE MODEL OF CONGENITAL CENTRAL HYPOVENTILATION SYNDROME

**DOI:** 10.1101/2021.02.15.431285

**Authors:** Amélia Madani, Gabriel Pitollat, Eléonore Sizun, Laura Cardoit, Maud Ringot, Thomas Bourgeois, Nelina Ramanantsoa, Christophe Delclaux, Stéphane Dauger, Marie-Pia d’Ortho, Muriel Thoby-Brisson, Jorge Gallego, Boris Matrot

**Author notes:** These authors contributed equally to this work (Amélia Madani, Gabriel Pitollat). **Corresponding author** Jorge Gallego, NeuroDiderot, INSERM, Hôpital Robert Debré, 48 Bd Sérurier, 75019 Paris, France. **Author contributions** J.G., B.M., S.D., C.D., M.T.B and M.P.O. conceived and designed the study, and prepared the first draft of the manuscript. B.M. designed and produced the device for apnea measurement and classification, T.B. validated this device and contributed to data analysis. A.M., E.S., M.R. acquired and analyzed respiratory measurements, N.R. generated the mutant mouse lines and supervised statistical analyses. G.P., L.C. and M.T.B., acquired and analyzed the histological and electrophysiological data. All authors contributed to the interpretation of the findings and critically revised the entire manuscript.

## Abstract

**Rationale:** Congenital Central Hypoventilation Syndrome (CCHS) is characterized by life-threatening sleep hypoventilation, and is caused by *PHOX2B* gene mutations, most frequently the *PHOX2B*^*27Ala*/+^ mutation, with patients requiring lifelong ventilatory support. It is unclear whether obstructive apneas are part of the syndrome.

**Objectives:** To determine whether *Phox2b*^*27Ala*/+^ mice, which present the main symptoms of CCHS and die within hours after birth, also express obstructive apneas, and to investigate potential underlying mechanisms.

**Methods:** Apneas were classified as central, obstructive or mixed by using a novel system combining pneumotachography and laser detection of abdominal movement immediately after birth. Several respiratory nuclei involved in airway patency were examined by immunohistochemistry and electrophysiology in brainstem-spinal cord preparation.

**Measurements and Main Results:** The median (interquartile range) of obstructive apnea frequency was 2.3/min (1.5-3.3) in *Phox2b*^*27Ala*/+^ pups versus 0.6/min (0.4-1.0) in wildtypes (*P* < 0.0001). Obstructive apnea duration was 2.7s (2.3-3.9) in *Phox2b*^*27Ala*/+^ pups versus 1.7s (1.1-1.9) in wildtypes (*P* < 0.0001). Central and mixed apneas presented similar, significant differences. In *Phox2b*^*27Ala*/+^ preparations, the hypoglossal nucleus had fewer (*P* < 0.05) and smaller (*P* < 0.01) neurons, compared to wildtypes. Importantly, coordination of phrenic and hypoglossal motor activities was disrupted, as evidenced by the longer and variable delay of hypoglossal with respect to phrenic activity onset (*P* < 0.001).

**Conclusions:** The *Phox2b*^*27Ala*/+^ mutation predisposed pups not only to hypoventilation and central apneas, but also to obstructive and mixed apneas, likely due to hypoglossal dysgenesis. These results thus demand attention towards obstructive events in infants with CCHS.

## Introduction

Congenital Central Hypoventilation Syndrome (CCHS) is an autosomal dominant disease whose incidence, although unknown precisely, ranges from 1/50.000 to 1/200.000 live births (1). CCHS is characterized by a loss of ventilatory chemosensitivity to hypercapnia and hypoxia, and life-threatening hypoventilation either exclusively during sleep or permanently in the most severe cases. The symptoms are generally noticed at birth or during early infancy. Patients typically require lifelong positive pressure ventilation, at least nocturnally, via tracheostomy during early development, and subsequently, through nasal or facial masks. CCHS is caused by heterozygous mutations in the paired-like homeobox 2B gene (*PHOX2B*) (2–4) that encodes a key transcription factor in the development of the autonomic nervous system (5). In 90% of cases, mutations are tri-nucleotide expansion mutations in *PHOX2B* leading to polyalanine expansions in the gene product (6). The most prevalent mutation is the 7-alanine expanded stretch (*PHOX2B*^*27Ala*/+^). The non-polyalanine mutations (missense, frameshift, nonsense and stop codon mutations) produce even more severe CCHS-related defects, including the need for continuous ventilatory support, Hirschsprung disease and tumors.

Central apneas have been commonly reported in CCHS patients and were assigned to a failure of the central respiratory drive from the brainstem respiratory centers (7, 8). Obstructive apneas were occasionally reported in case or small-group studies (9–17) and were considered to require management in 50% of patients under one year old (18). However, with rare exceptions (19), the study of apneas in CCHS was conducted before the discovery of the disease-causing gene (2, 4), and possibly included heterogeneous patient groups. In current practice, obstructive apneas if occurring, may remain unnoticed due to tracheostomy and/or mechanical ventilation, which is generally used in a bi-level positive pressure mode (20). Thus, the question of whether obstructive disorders are part of CCHS remains unanswered, which may confound ongoing search for effective pharmacological treatments.

Here, we analyzed whether the *Phox2b*^*27Ala*/+^ mutation causes obstructive apneas in mice. Previous studies showed that *Phox2b*^*27Ala*/+^ mice present main CCHS symptoms and die from respiratory arrest within hours after birth (21). Furthermore, the retrotrapezoid nucleus, a cluster of central chemoreceptors providing the major excitatory drive to breathing, is severely depleted (22, 23). Accordingly, apneas in *Phox2b*^*27Ala*/+^ mice were assigned to retrotrapezoid nucleus agenesis and considered to be central in origin. However, the *Phox2b*^*27Ala*/+^ mutation may also potentiate upper airway obstruction because *Phox2b*-expressing regions containing hypoglossal premotor neurons may be functionally impaired (24). On this basis, we hypothesized that *Phox2b*^*27Ala*/+^ pups may express not only central, but also obstructive and mixed apneas. To test this idea, we developed an original method that enables classifying apneas as central, obstructive or mixed in newborn mice. We then analyzed apneas in *Phox2b*^*27Ala*/+^ pups and sought their neuronal basis by examining late stage fetal *Phox2b*^*27Ala*/+^ embryos.

## Methods

### Mice

We generated pups with the constitutive *Phox2b*^*27Ala*/+^ mutation (21), hereafter termed *Phox2b*^*27Ala*/+^ pups, and pups with the conditional, tissue-specific *Phox2b*^*27Ala*/+^ mutation targeted to the retrotrapezoid nucleus (25), hereafter termed *Phox2b*^*27Alacond*/+^ pups. See online data supplement for the ethical approval and details on mutant generation and genotyping.

### Pneumotachography

Breathing variables were measured using a pneumotachograph and a facemask combined into one single 3D-printed component (termed “pneumotachometer”; Figures 1A and 1B, see details in the online data supplement). The pneumotachometer was calibrated before each monitoring session. The flow signal was integrated for breath-by-breath measurement of tidal volume using a custom-made software. We measured breathing frequency (f_R_, number of breaths/min), tidal volume (divided by body weight, V_T_, mL/g), ventilation (V_E_, calculated as f_R_ x V_T_, mL/g/min) and apneas (detailed below). Breathing variables and apneas were analyzed over activity-free periods based on video recordings (Logitech C920, Lausanne, Switzerland), which were considered as corresponding to behavioral sleep.

**Figure 1.**
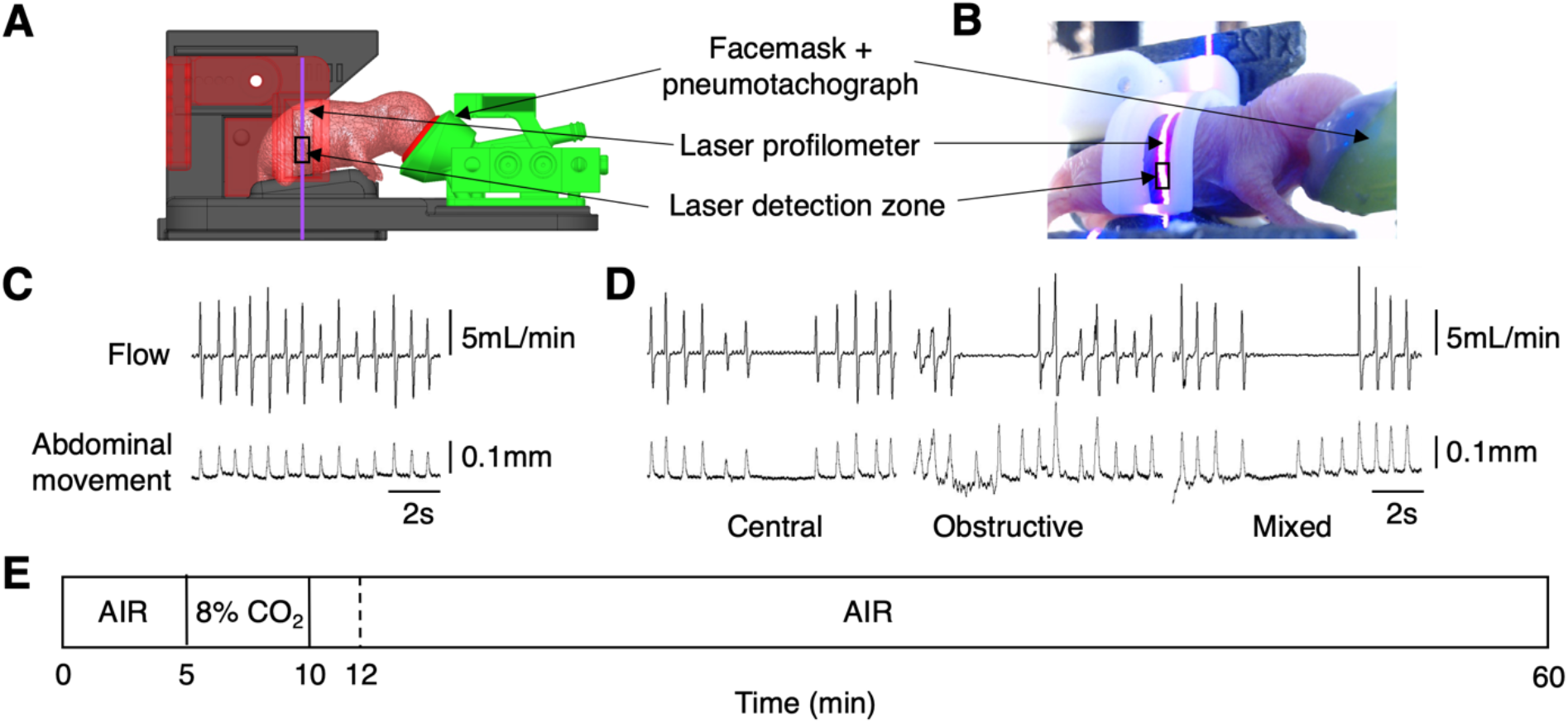
Experimental setup for detection and classification of central, obstructive and mixed apneas in newborn mice. (*A*) Technical drawing of the ‘pneumotachometer’ incorporating a Fleisch pneumotachograph and a facial mask that was fixed on the pup’s snout. Air flow through the mask (20 mL/min^−1^) prevented rebreathing. A laser profilometer detected abdominal movements within a 2 mm x 0.5 mm zone (black rectangle). (*B*) Image from video recording showing the facemask and the laser beam. The entire setup was placed in a thermoregulated chamber (not shown). (*C*) Individual traces of flow signal from the pneumotachograph (upper trace) and movement monitored by laser profilometry (lower trace) from a pup. (*D*) Examples of central (no flow, no movement), obstructive (no flow, abdominal movement) and mixed (no flow, abdominal movement at the end of apnea) in a *Phox2b*^*27Ala*/+^ pup. (*E*) Experimental protocol. After 5 min of air breathing, each pup was exposed to 8% CO_2_, and then back to air for 50 min recording. Apneas were classified as central, obstructive or mixed from minute 12 to minute 60.

### Laser detection of respiratory efforts

Abdominal movements were measured using a laser profilometer pointing radially at the lateral abdominal wall (Figures 1A and 1B) to detect respiratory efforts during obstructive events (see details in the online data supplement).

### Apnea detection and classification

Apneas were classified as central, obstructive or mixed by adjusting the American Academy of Sleep Medicine recommendations (26) to newborn mice. An apnea was characterized by a > 90% decrement in peak inspiratory airflow for a duration of at least 0.90 s, which corresponds approximately to two breaths in neonatal mice. The classification into central, obstructive, or mixed apneas was performed by visual inspection of pneumotachometer and laser profilometer signals. An apnea was considered as central when no respiratory effort was present throughout the apnea, as obstructive when associated with continuous respiratory effort, and as mixed when an initial period lacking abdominal movement was followed by resumption of respiratory effort (Figure 1D).

### Electrophysiology

Electrophysiological recordings of rhythmically organized breathing-related activities were performed on isolated brainstem preparations harvested the day before birth (embryonic day 18.5) as previously described (27) (see details in the online data supplement).

### Protocol

Each pup was left with its mother for 15-20 min until the end of maternal care. The pup was then sexed, weighed and attached to the pneumotachometer inside a thermoregulated chamber (32°C). The snout was sealed into the facemask with polyether adhesive (Impregum, 3M, Saint Paul, MN, USA). Recordings were started within 100 min following delivery. Breathing variables were recorded for 60 min (Figure 1E) and extended beyond this period in *Phox2b*^*27Ala*/+^ pups until terminal apnea to analyze gasping.

### Statistics

For normally distributed data, we used two tailed t-tests and two-way ANOVAs, followed by a Bonferroni’s multiple comparisons test where appropriate. Non-normally distributed data were analyzed using the Mann-Whitney test, followed by Dunn’s post-hoc multiple comparisons test. Sex had no statistically significant effects in any analysis and will not be mentioned further. All analyses were performed using Prism 9 (GraphPad). *P* values < 0.05 were considered significant.

## Results

### Baseline breathing and apnea time in *Phox2b*^*27Ala*/+^ pups

The breathing pattern of *Phox2b*^*27Ala*/+^ pups was typified by significantly reduced ventilation and increased apnea total time (irrespective of apnea type), compared to wildtype pups (Table 1), thereby confirming previous plethysmographic data (21). The lower V_E_ of *Phox2b*^*27Ala*/+^ pups was mainly due to a lower f_R_. The V_E_-response to hypercapnia was abolished in *Phox2b*^*27Ala*/+^ pups, as previously reported (21) despite a small significant V_T_-response (see Figures E2A and E2B in the online data supplement).

**Table 1.** Baseline breathing variables in constitutive *Phox2b*^*27Ala*/+^ newborn mice and wildtype controls. Data were collected from minute 12 to minute 60 of the protocol. Movement time was expressed in seconds per minute of recording time. f_R_: breathing frequency; V_T_: tidal volume; V_E_: mean ventilation. Apnea time was expressed in seconds per minute of movement-free time (corresponding to sleep), and calculated irrespective of apnea type. *P* values: t-test comparison between mutants and wildtypes. Values are means ± SD.

| Variable | Wildtype<br>n = 48 | *Phox2b*<sup>27Ala/+</sup><br>n = 36 | *P* value |
| --- | --- | --- | --- |
| Weight (g) | 1.33 ± 0.09 | 1.28 ± 0.1 | 0.007 |
| Movement time (s/min) | 24 ± 7 | 15 ± 8 | < 0.001 |
| f<sub>R</sub> (breaths/min) | 98 ± 18 | 46 ± 19 | < 0.001 |
| V<sub>T</sub> (mL/g x 10<sup>-3</sup>) | 6.7 ± 0.9 | 6.3 ± 1.4 | 0.104 |
| V<sub>E</sub> (mL/min/g) | 0.65 ± 0.13 | 0.29 ± 0.16 | < 0.001 |
| Apnea time (s/min) | 7.22 ± 4.60 | 26.67 ± 12.79 | < 0.001 |

### Central, obstructive and mixed apneas in *Phox2b*^*27Ala*/+^ pups

Compared to wildtypes, the *Phox2b*^*27Ala*/+^ pups expressed significantly more severe central, obstructive and mixed apneas, in terms of mean duration, frequency and total time (Figure 2). The total time for central apneas was longer (Figure 2A) due to both higher apnea occurrence frequencies (Figure 2B), and their longer mean durations (Figure 2C). This result confirmed and extended previous claims that the *Phox2b*^*27Ala*/+^ mutation causes central apneas (21), but here, based on formal apnea classification. More importantly, the total time for obstructive apneas, which hitherto has never been specifically investigated, was longer in *Phox2b*^*27Ala*/+^ pups compared to wildtypes (Figure 2A), and this was due to a higher apnea frequency (Figure 2B), and longer mean duration (Figure 2C). A comparison of mixed apneas between *Phox2b*^*27Ala*/+^ pups and wildtypes yielded the same statistical results as for central and obstructive apneas (Figures 2A-C)

**Figure 2.**
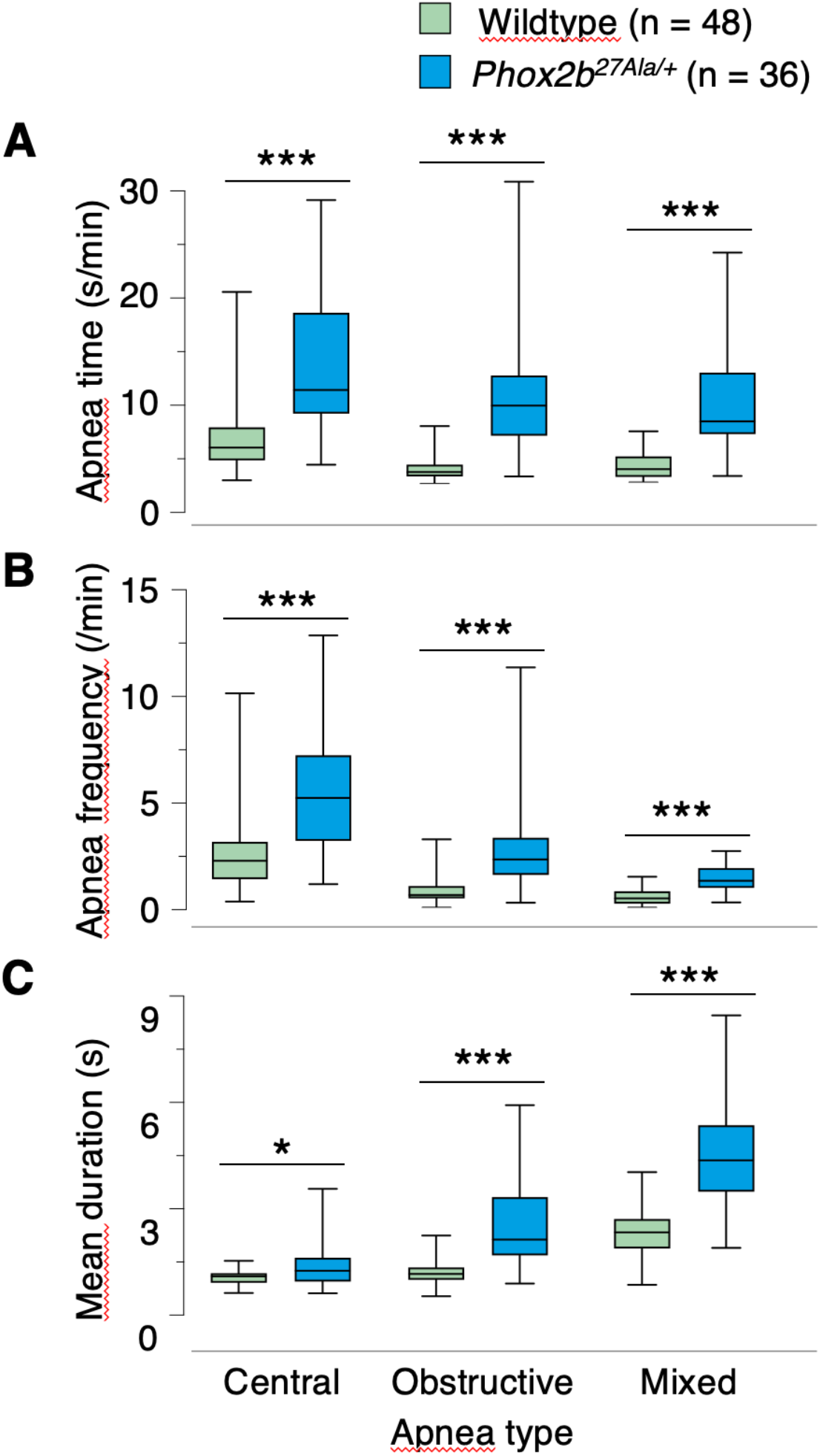
The *Phox2b*^*27Ala*/+^ mutation enhanced central, obstructive and mixed apneas in newborn pups compared to wildtypes. Apneas were quantified according to Total time (*A*) calculated as the ratio of total apnea durations divided by recording time after exclusion of activity periods, Apnea frequency (*B*), and Apnea mean duration (*C*). Undetermined apneas represented less than 2% of total apnea time in either group and were not considered. The high interindividual variability in the apnea characteristics of mutants reflected their variable lifetime, which ranged from a few minutes to 24 hours. Lines and whiskers in boxplots represent medians, interquartile ranges, and minimum and maximum values. \**P* = 0.0129, \*\*\**P* < 0.0001 indicate significant differences.

Finally, the proportion of obstructive apneas (with respect to total apnea number) was higher in *Phox2b*^*27Ala*/+^ pups, compared to wildtype pups (29% ± 14 *vs* 23% ± 13, respectively, *P* < 0.05), and the proportion of time spent in obstructive apneas (with respect to total apnea time) was also longer in *Phox2b*^*27Ala*/+^ pups (31% ± 14 *vs* 22% ± 13, respectively, *P* < 0.001).

### Obstructed gasping in *Phox2b*^*27Ala*/+^ newborn mice

We next examined the possibility that airway obstruction contributed to the death of *Phox2b*^*27Ala*/+^ pups in a subsample of mutant pups (n = 45) in which respiratory monitoring was maintained until respiratory arrest. The occurrence of gasps is illustrated in the recording sequences from a single pup in Figure 3. The initial period during which breathing was variably affected (Figures 3A and 3B), was followed by a period of disorganized breathing interspersed with gasps, which were characterized by large and brief respiratory efforts accompanied by large inspirations (Figure 3C). Some of these large abdominal movements were not associated with inspiration, indicating obstructed gasps (Figure 3D). In the 3-min period preceding respiratory arrest, 25 out of the 45 (55%) pups produced such obstructed gasps.

**Figure 3.**
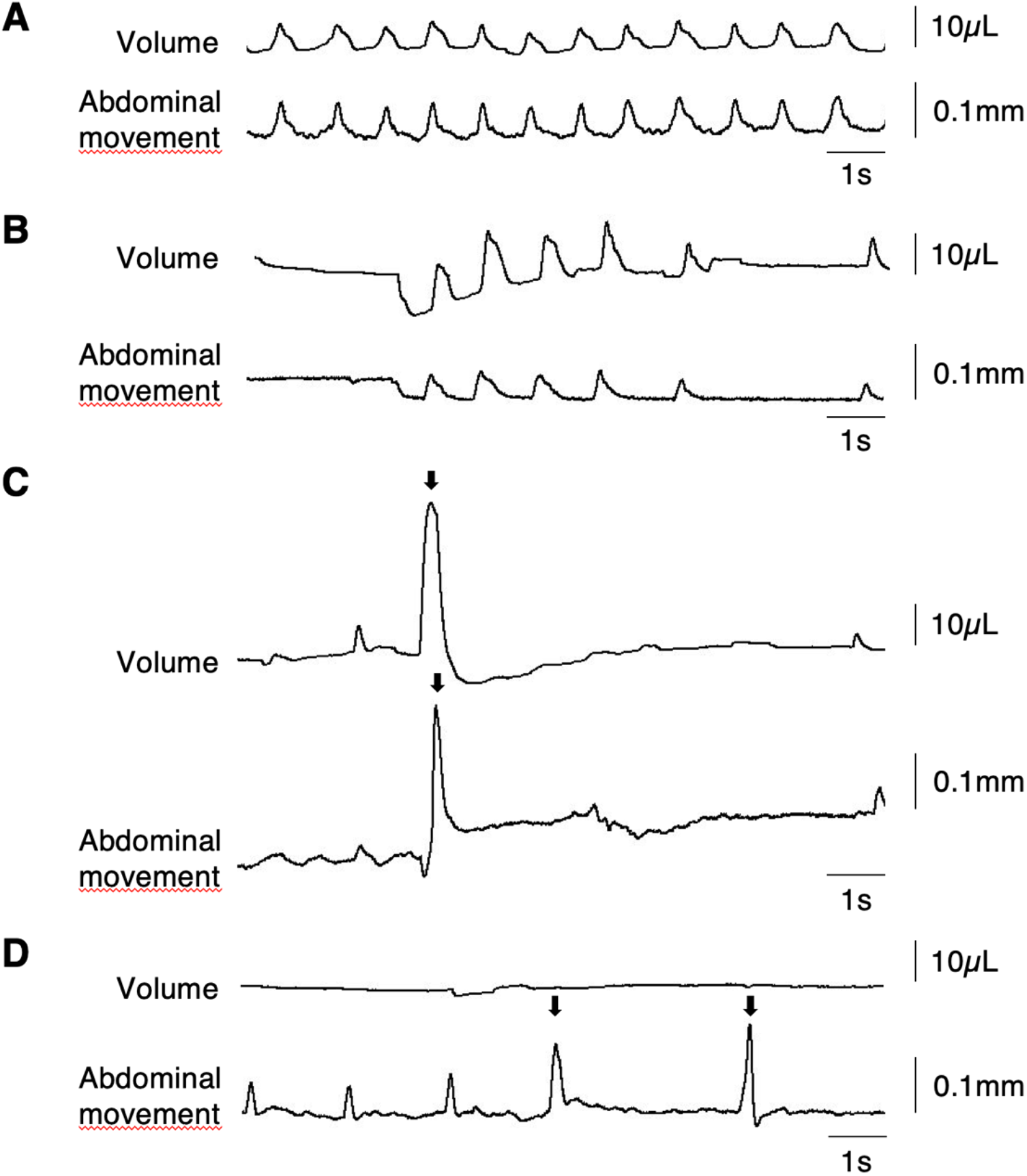
Examples of unobstructed and obstructed terminal gasps in a *Phox2b*^*27Ala*/+^ pup. (*A*) Slow, regular breathing. (*B*) Breathing interspersed with apneas. (*C*) Irregular breathing with apneas and an unobstructed gasp accompanied by an abdominal expansion (arrows). (*D*) Long obstructive apnea followed by two obstructed gasps (arrows). The four excerpts in (*A-D*) were recorded in the same *Phox2b*^*27Ala*/+^ pup after 3h02, 4h15, 4h22 and 4h52 of recording, respectively.

### Implication of retrotrapezoid neuronal loss in apneas

We asked whether the obstructive pattern of *Phox2b*^*27Ala*/+^ pups was due to an agenesis of the brainstem retrotrapezoid nucleus, which is the unique neuroanatomical defect reported to date in *Phox2b*^*27Ala*/+^ pups (21). To this end, we replicated the main study in conditional knock-in, tissue-specific mice, in which the *Phox2b*^*27Ala*/+^ mutation was targeted to the retrotrapezoid nucleus region. These mutants, hereafter designated *Phox2b*^*27Alacond*/+^ pups, previously analyzed at postnatal day two, survived normally without apneas, despite the agenesis of retrotrapezoid nucleus and the consequent loss of CO_2_-chemosensitivity (25). Here, we found that within 100 min after birth, *Phox2b*^*27Alacond*/+^ pups had a lower V_E_, mainly due to lower f_R_ values, and longer total apnea times (irrespective of apnea type) compared to wildtypes (Table 2). Moreover, the ventilatory response to CO_2_ was almost abolished in *Phox2b*^*27Alacond*/+^ pups (see Figures E3A and E3B in the online data supplement) consistent with a retrotrapezoid neuronal loss. The sole loss of retrotrapezoid neurons in *Phox2b*^*27Alacond*/+^ pups not only increased the frequency and total time of central apneas, but it also increased the mean duration of obstructive apneas compared to wildtypes (Figure 4A-C). No statistical comparison was made between *Phox2b*^*27Ala*/+^ and *Phox2b*^*27Alacond*/+^ pups, since these data were collected from two different studies, but an approximate assessment of apnea characteristics in the two mutants clearly indicated that the apneic phenotype was much more severe in *Phox2b*^*27Ala*/+^ than in *Phox2b*^*27Alacond*/+^ pups. Thus, the exclusive loss of neurons in the retrotrapezoid nucleus promoted central apneas and aggravated obstructive apnea in *Phox2b*^*27Alacond*/+^ pups, but it could not account for the dramatic apneic phenotype of *Phox2b*^*27Ala*/+^ pups.

**Table 2.** Baseline breathing variables in conditional tissue-specific *Phox2b*^*27Alacond*/+^ newborn mice and wildtype controls. Data were collected from minute 12 to minute 60 of the protocol. Movement time was expressed in seconds per minute of recording time. f_R_: breathing frequency; V_T_: tidal volume; V_E_: mean ventilation. Apnea time (in seconds per minute of movement-free time) was calculated irrespective of apnea type. *P* values: t-test comparisons between mutants and wildtypes. Values are means ± SD.

| Variable | Wildtype<br>n = 62 | *Phox2b*<sup>27Alacond/+</sup><br>n = 58 | *P* value |
| --- | --- | --- | --- |
| Weight (g) | 1.31 ± 0.12 | 1.35 ± 0.10 | 0.0198 |
| Movement time (s/min) | 27 ± 10 | 24 ± 8 | 0.049 |
| f<sub>R</sub> (breaths/min) | 97 ± 28 | 79 ± 24 | < 0.001 |
| V<sub>T</sub> (mL/g x 10<sup>-3</sup>) | 6.6 ± 0.9 | 6.3 ± 0.8 | 0.079 |
| V<sub>E</sub> (mL/min/g) | 0.64 ± 0.19 | 0.49 ± 0.14 | < 0.001 |
| Apnea time (s/min) | 8.08 ± 5.92 | 11.34 ± 8.62 | 0.017 |

**Figure 4.**
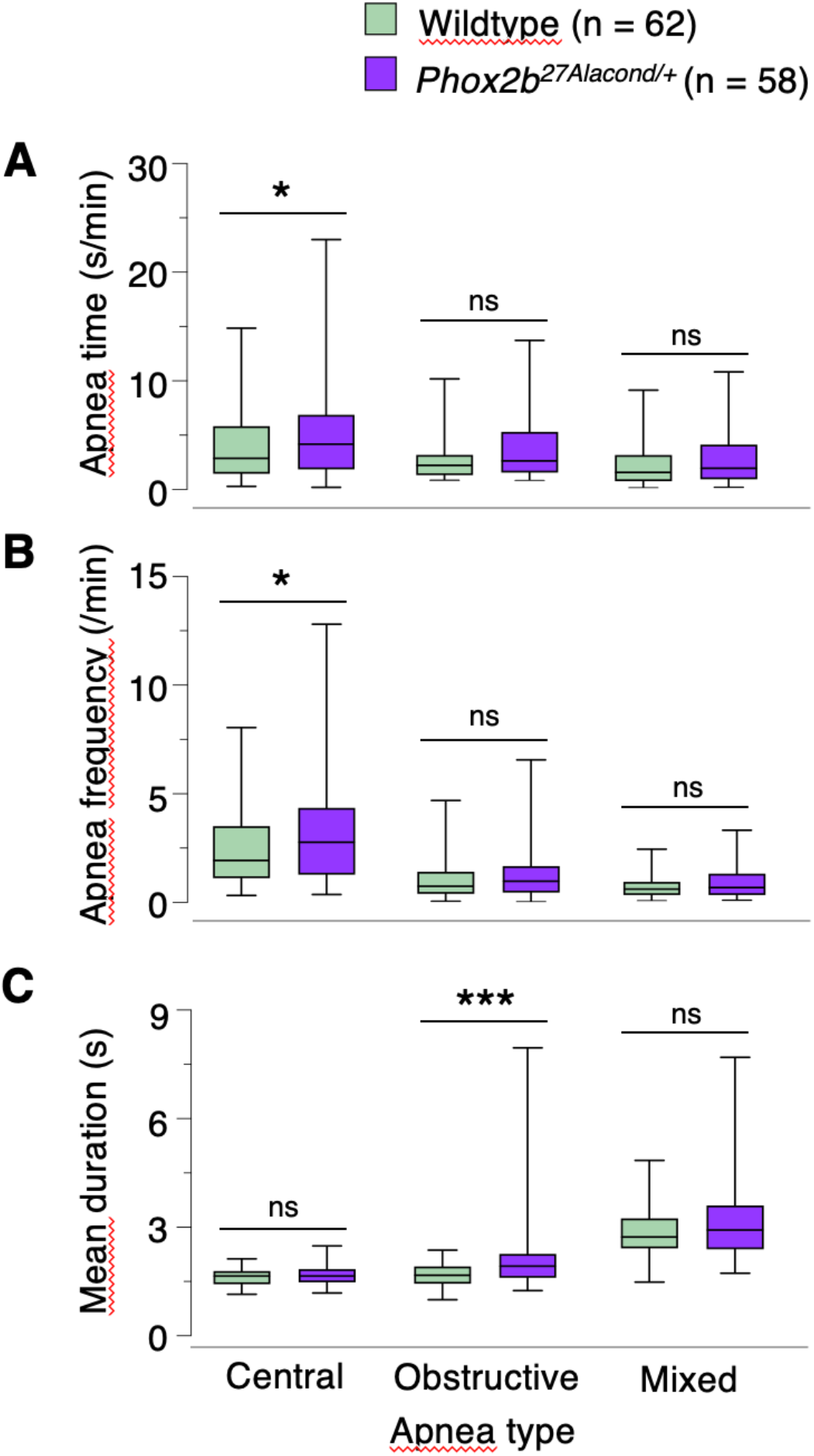
The conditional, tissue-specific *Phox2b*^*27Alacond*/+^ mutation enhanced central and obstructive apnea expression compared to wildtypes. Apneas were quantified by Total time (*A*) calculated as the ratio of total apnea durations divided by the recording time after the exclusion of activity periods, Apnea frequency (*B*), and Mean duration (*C*). Lines and whiskers in boxplots represent medians, the interquartile ranges, and the minimum and maximum values. Compared with *Phox2b*^*27Ala*/+^ pups (Figure 2), the apneic phenotype of *Phox2b*^*27Alacond*/+^ was much less severe. \**P* < 0.05, \*\*\**P* < 0.001 indicate significant differences, ns indicates no statistical difference.

### Dysgenesis and dysfunction of the hypoglossal nucleus in newborn *Phox2b*^*27Ala*/+^ mice

Because obstructive and mixed apneas in *Phox2b*^*27Ala*/+^ pups were not mainly attributable to retrotrapezoid nucleus agenesis, we examined the hypoglossal nucleus, which is the main group of motoneurons involved in the control of the upper airway muscles (28). In *Phox2b*^*27Ala*/+^ pups, the hypoglossal nucleus displayed an abnormal anatomy (Figures 5A and 5B) characterized by a smaller perimeter (Figure 5C), a reduced surface (Figure 5D), and fewer neurons (Figure 5E), despite having an apparently normal rostrocaudal extension (Figure 5F) and neuroanatomical location (Figure 5G).

**Figure 5.**
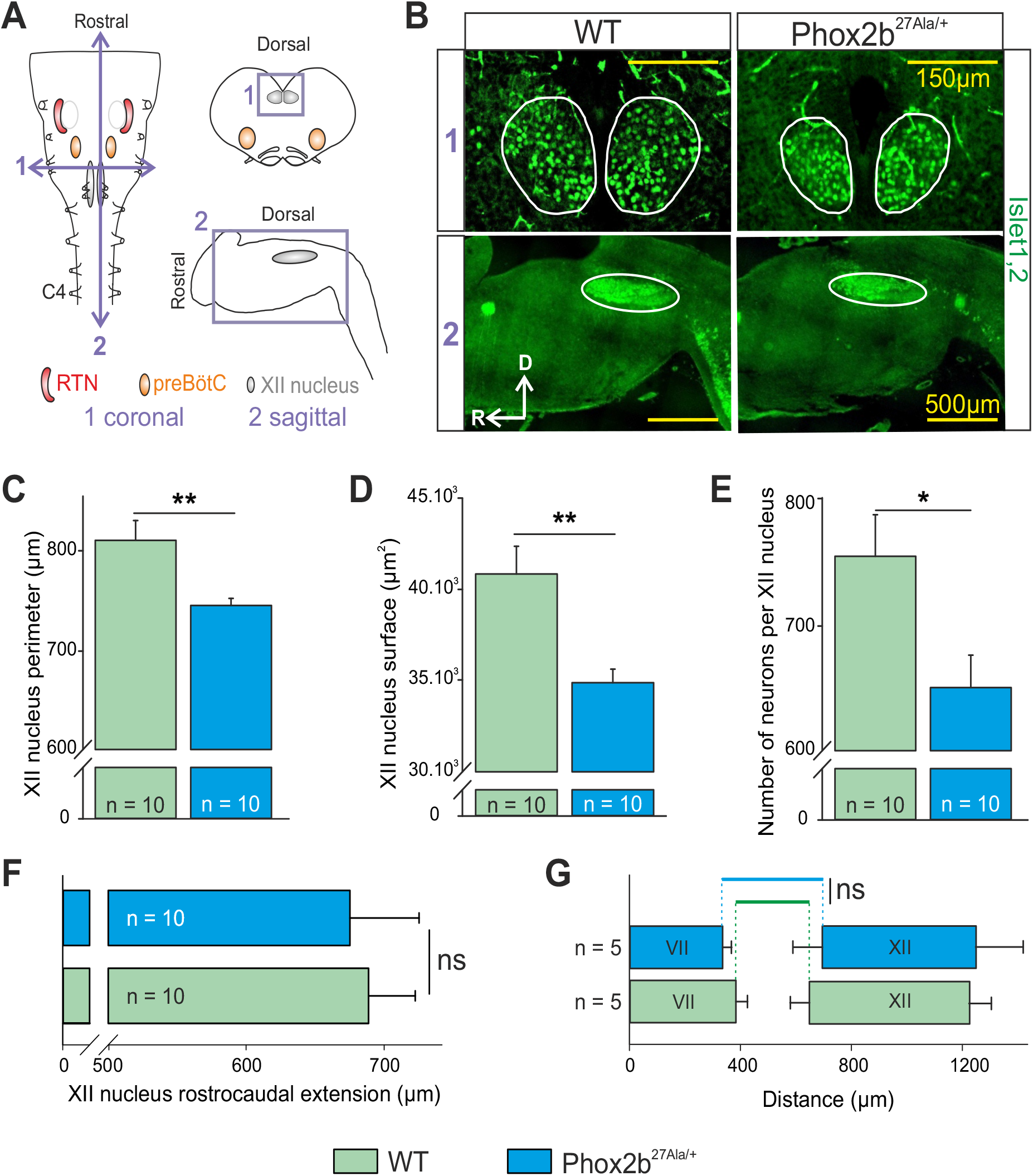
Abnormal anatomy of the hypoglossal nucleus in the *Phox2b*^*27Ala*/+^ mutant. (*A*) Schematic representations of an isolated brainstem preparation (left), a coronal brainstem slice (right, top) and a sagittal brainstem slice (right, bottom) with the corresponding position of the retrotrapezoid nucleus (RTN; red shading), the preBötzinger complex (preBötC; orange shading) and the hypoglossal (XII) nucleus (gray shading). The double-headed violet arrows indicate the coronal (1) and sagittal (2) plans of section. The violet rectangles delimit the areas shown in B. (*B*) Immunostainings against Islet1,2 (green) in 30 μm thick brainstem coronal (top) and sagittal (bottom) slices obtained from embryonic day 18.5 WT and *Phox2b*^*27Ala*/+^ embryos. White lines delimit the XII nuclei. (*C-G*) Histograms showing overall mean values (±SEM) for the XII nucleus perimeter (*C*) and surface (*D*), the number of neurons per XII nucleus (*E*), the rostro-caudal extension distance (*F*) and the position relative to the caudal end of the VII nucleus (*G*) of the XII nucleus measured from WT (green) and *Phox2b*^*27Ala*/+^ (blue) embryos. The number of preparations analyzed is indicated in each bar graph. \**P* < 0.05, \*\**P* < 0.01 indicate significant differences (t-test), ns indicates no statistical difference. In the mutant, the XII nucleus is smaller, contains less neurons but with the same rostro-caudal extension and position relative to the VII nucleus. C4: phrenic rootlet, RTN: retrotrapezoïd nucleus, preBötC: preBötzinger complex, VII: facial nucleus, XII: hypoglossal nucleus, WT: wildtype.

To examine whether the function of the hypoglossal nucleus is affected by the *Phox2b*^*27Ala*/+^ mutation, we recorded spontaneous hypoglossal and phrenic nerve activity in embryonic (E)18.5 isolated brainstem-spinal cord preparations that retain the ability to express respiratory-related phrenic and hypoglossal activities *in vitro* (27). Electrophysiological monitoring of the hypoglossal nucleus revealed significant functional disorders in *Phox2b*^*27Ala*/+^ preparations (Figure 6A). Specifically, burst frequency was significantly reduced in *Phox2b*^*27Ala*/+^ preparations compared to wildtypes (Figure 6B), in line with their reduced breathing frequency *in vivo,* and the rise time of burst intensity was also reduced, but without significantly affecting burst duration (Figure 6C-E). Most importantly, the combined analysis of hypoglossal and phrenic nerve activities showed a marked loss of coordination between these two neural activities. Specifically, the mutant preparations presented a longer and more variable delay of hypoglossal activity onset with respect to that of the phrenic nerve, which contrasted with the near coincident activities of wildtype preparations (Figures 6F and 6G). The large intra-individual variability of the delay between hypoglossal and phrenic burst onsets (Figure 6F) was observed in all *Phox2b*^*27Ala*/+^ preparations, and further pointed to an impairment of the coordination between these activities.

**Figure 6.**
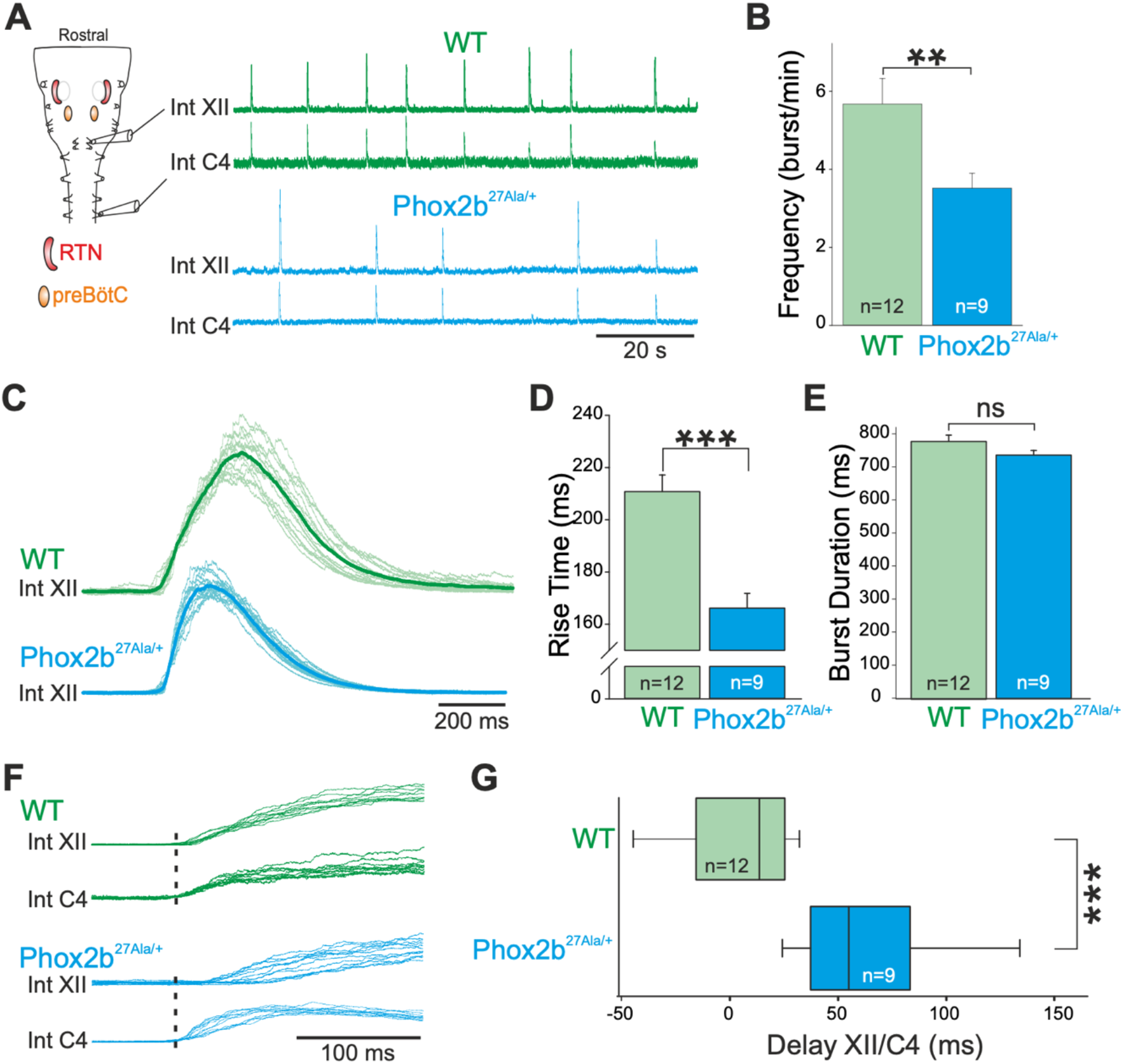
The hypoglossal nerve is functional but the shape of its bursts and their coordination with phrenic nerve activity are altered in the *Phox2b*^*27Ala*/+^ mutant. (*A*) Left, Schematic representation of an isolated brainstem preparation with the arrangement for simultaneous electrophysiological recording of hypoglossal and phrenic nerve activities. Right, Representative rectified and integrated (Int) simultaneous recordings from hypoglossal (XII) and phrenic (C4) ventral roots in *in vitro* preparations obtained from wildtype (WT, green traces) and a mutant (*Phox2b*^*27Ala*/+^, blue traces) mice. (*B*) Histograms showing mean XII burst frequency in 12 WT (green) and 9 *Phox2b*^*27Ala*/+^ (blue) preparations. (*C*) Superimposed XII bursts and average traces (bold line) from representative recordings of a WT (n = 10 bursts, green) and a *Phox2b*^*27Ala*/+^ (n = 12 bursts, blue) preparation. (*D*) and (*E*) Histograms showing mean rise times and durations of XII bursts in WT (n = 120 events, green) and *Phox2b*^*27Ala*/+^ (n = 90 events, blue) recordings. (*F*) Superimposition of spontaneous hypoglossal (top, n = 10 bursts) and phrenic (bottom, n = 12 bursts) motor bursts recorded from a WT (green) and a *Phox2b*^*27Ala*/+^ (blue) preparation. The dotted lines indicate the mean onset of the phrenic bursts. Note that the coordination between phrenic and hypoglossal burst onsets is affected in the *Phox2b*^*27Ala*/+^ preparations. (*G*) Boxplots with whiskers (min to max) representing the mean delay (± SEM) between phrenic and hypoglossal bursts. Hypoglossal bursts were initiated later than phrenic bursts in the *Phox2b*^*27Ala*/+^ preparations and over a wider range of values. \*\**P* < 0.01, \*\*\**P* < 0.001 indicate significant differences (Mann-Whitney t-test), ns indicates no statistical difference. C4: phrenic rootlet, RTN: retrotrapezoïd nucleus, preBötC: preBötzinger complex, XII: hypoglossal nucleus, WT: wildtype.

## Discussion

Here, we show that, compared to wildtypes, newborn mice bearing the *Phox2b*^*27Ala*/+^ constitutive mutation display an abnormally high degree of not only central but also obstructive and mixed apneas, with the latter tending to be longer than central apneas. Furthermore, terminal apneas were frequently preceded by obstructed gasps, suggesting they play a causal or aggravating role in some pups. The obstructive phenotype of *Phox2b*^*27Ala*/+^ is ascribed at least partly to dysgenesis and dysfunction of the hypoglossal nucleus, which is critical to the maintenance of upper airway patency.

### Methodology for apnea detection and classification in newborn mice

Our study is the first to discriminate apneas according to type (central, obstructive and mixed) in newborn mice, thanks to an original method that combines a built-in precision pneumotachometer and an ultra-sensitive laser profilometer. Previous attempts to classify apneas in newborn rodents were developed in 7-day old rats using electromyographic electrodes implanted into respiratory muscles under anesthesia (29). This approach was not applicable in the present study due the short lifetime (70% died within three hours after delivery) and high vulnerability to anesthesia of *Phox2b*^*27Ala*/+^ mutants (30). Despite our efforts to adjust the facemask to the pups’ morphology, the orofacial contact with the mask may have influenced sleep-wake states and breathing. To avoid this effect, we systematically discarded all periods of gross body movements monitored in video recordings to retain behavioral sleep periods only. However, we cannot completely exclude the possibility that some periods of quiet wakefulness were included in our apnea analysis, irrespective of genotype.

A further limitation of our study is that we did not attempt to discriminate sleep states, because, at early postnatal stage examined, this would have required the invasive measure of nuchal EMG (31). Thus, the sleep state-dependence pattern of apnea characteristics was not investigated. Previous studies on 5-day old mice have indicated that active sleep occupies approximately 80% of sleep time, in line with human data (31), thereby suggesting that the apneic patterns described here mainly reflect those that developed during active sleep. Despite these limitations, the present methodology allows the classification of apneas in an infant mouse model of CCHS, with potential applications to other rodent models of sleep disordered breathing in infants.

### Apnea profiles in *Phox2b*^*27Ala*/+^ newborn mice

As noted above, apneas in *Phox2b*^*27Ala*/+^ pups were previously ascribed to loss of CO_2_-driven excitatory input to respiratory rhythm generator by the retrotrapezoid nucleus. All apneas were therefore considered as central. The main result of the present study is that the *Phox2b*^*27Ala*/+^ mutation not only leads to central apneas, but also to obstructive and mixed apneas. On one hand, central apneas were found to be more frequent than obstructive and mixed apneas, and consequently could be expected to produce more numerous hypoxia-reoxygenation sequences, which are considered to be the main cause of apnea-related morbidity. On the other hand, obstructive and mixed apneas were significantly longer than central apneas, and thus would potentially be associated with larger decreases in heart rate and oxygen saturation (32). Furthermore, the co-occurrence of obstructive events and terminal apnea suggested that obstructive events were responsible for precipitating terminal apnea and death in many *Phox2b*^*27Ala*/+^ pups. Taken together, these considerations support the conclusion that obstructive events caused by the *Phox2b*^*27Ala*/+^ mutation may seriously affect morbidity and mortality in the neonatal period.

### Neural basis for obstructive apneas in *Phox2b*^*27Ala*/+^ newborn mice

The hypoglossal motor nucleus innervates the muscles of the tongue, and therefore withdrawal of an excitatory drive of hypoglossal motoneurons is a common cause of obstructive apneas (28). The dysgenesis of the hypoglossal nucleus found in the present study is a novel result. In previous studies, neuroanatomical defects were sought primarily in *Phox2b*-expressing structures, leading to the discovery of retrotrapezoid nucleus agenesis (21), without consideration of the hypoglossal nucleus, which does not depend on *Phox2b* for its development (24). Our finding of a dysgenesis of the hypoglossal nucleus thus showed that the *Phox2b*^*27Ala*/+^ mutation elicits non-cell-autonomous mechanisms affecting respiratory networks. To date, such mechanisms were only reported in newborn mice with a non-polyalanine repeat mutation (the frameshift *Phox2BΔ8* mutation), which expressed a dysfunctional preBötzinger complex, a non-*Phox2b* dependent brainstem site that is responsible for respiratory rhythmogenesis (33). Taken together, therefore, these findings widen the range of possible respiratory control disorders caused by *Phox2b* mutations.

The neuroanatomical defects of the hypoglossal nucleus were associated with functional defects, the most significant being the uncoupling of hypoglossal and phrenic nerve activities. In normal physiological conditions, the hypoglossal nucleus tends to burst synchronously with the phrenic nerve in order to exert tonic input to the upper airway dilator muscle motor neurons, thereby preventing airway collapse during inspiratory depression (34, 35). Here, the onset of hypoglossal discharge was found to be much less well coordinated with diaphragmatic activity in *Phox2b*^*27Ala*/+^ pups, in contrast to the wildtype. This in turn suggested that, *in vivo*, the inspiratory negative pressure in *Phox2b*^*27Ala*/+^ pups is not counteracted by a tonic activation of the upper airway muscles, at least during the initial part of inspiration, thus increasing the propensity for obstructive events. In addition to the likely impact of hypoglossal dysfunction on obstructive events, we also showed that the loss of neurons solely in the retrotrapezoid nucleus was associated with longer obstructive apneas (as shown in *Phox2b*^*27Alacond*/+^ pups). This result thus provides *in vivo* support for the notion that the retrotrapezoid nucleus makes a significant contribution to the control of genioglossus muscle activity (36, 37). Thus, the outcome of hypoglossal and retrotrapezoidal defects acting in combination are likely to further aggravate obstructive apneas in *Phox2b*^*27Ala*/+^ pups. However, we cannot exclude the possibility that hypoglossal activity is also affected by a dysfunction of other brain regions containing hypoglossal premotor neurons expressing *Phox2b* and possibly also affected by the mutation (24).

### Clinical relevance

The high incidence of not only central, but also obstructive and mixed apneas in *Phox2b*^*27Ala*/+^ newborn mice suggested that airway patency disorders also occur in infants with CCHS, but remain undetected due to permanent tracheostomy, and/or the use of positive pressure ventilation. Furthermore, our findings suggest that, if observed, obstructive events should not rule out a diagnosis of CCHS. This study also indicates that obstructive events in patients with CCHS need special attention in the perspective of future treatment research, which to date, has been mainly oriented toward the employment of respiratory stimulants (38–40). Finally, our data shed new light on the neurobiology and neurodevelopment of pharyngeal muscle control, with a potential translation towards pharmacotherapeutic treatment that targets pharyngeal patency and obstructive sleep apnea syndromes.

## Conclusion

We demonstrate that the *Phox2b*^*27Ala*/+^ mutation, which is the most frequent cause of CCHS, leads to a combination of central, obstructive and mixed apneas in newborn mice. These obstructive disorders are attributable at least in part, to a dysfunction of the hypoglossal nucleus, a non-*Phox2b* expressing structure, together with a disruption of input from the *Phox2b-* expressing retrotrapezoid nucleus. These results broaden the spectrum of neuroanatomical and neurofunctional disorders caused by *Phox2b* mutations, and point to the clinical importance of detecting obstructive events in infants with CCHS.

## ACKNOWLEDGEMENTS

We are deeply indebted to Dr. Florian Brunet, Sebastien Lanteri and Victoria Jean-Baptiste Sérichard for their contribution in developing the apnea detection device, to Fabien Duverger for his contribution to the pneumotachometer design, to Dr. Chih-Wei Hsu and Dr. Mary Dickinson for providing 3D scans of newborn mice, to Charline Riaud and Aurélie Hayotte for experimental assistance and to Dr. John Simmers for English editing.

