## Supplementary material for "OBSTRUCTIVE APNEAS IN A MOUSE MODEL OF CONGENITAL CENTRAL HYPOVENTILATION SYNDROME": ONLINE DATA SUPPLEMENT

### Methods

#### Ethical approval

All experimental protocols were approved by local (Bichat-Robert-Debré and University of Bordeaux) and national ethics committees (Ministère de l'Enseignement Supérieur et de la Recherche – Direction Générale pour la Recherche et l'Innovation, Ethical approval # 1278). The wildtype pups and conditional mutants were sacrificed by decapitation upon completion of the physiological recordings.

#### Mice

Two mutant lines were generated: the constitutive *Phox2b*$^{27Ala/+}$ mutant (1), designated as *Phox2b*$^{27Ala/+}$ pups, and the conditional, tissue-specific *Phox2b*$^{27Ala/+}$ mutation targeted to the retrotrapezoid nucleus (2), designated as *Phox2b*$^{27Alacond/+}$ pups. Instead of generating these mutants from chimeric male mice as previously described (1), we crossed *Pgk::Cre* male mice with floxed *Phox2b*$^{27Ala/27Ala}$ females (2). To restrain the mutation to the RTN structure, we crossed *Krox20*$^{Cre/+}$ male mice with floxed *Phox2b*$^{27Ala/27Ala}$ females (2). Genotyping was done on tail DNA using the following primers: (i) to detect the presence of the +7Ala allele GCCCACAGTGCCTCTTAACT and CTCTTAAACGGGCGTCTCAC yielding bands of 330 pb for the wild-type, of 474pb for the mutated and of 380kb for the recombined alleles, (ii) to detect cre AAATTTGCCTGCATTACCG and ATGTTTAGCTGGCCCAAATG yielding a band of 250 pb.

#### Pneumotachometer

The facemask was designed using SolidEdge (Siemens PLM Software, Plano, TX, USA) after 3D scans of the mouse body on the day of birth (courtesy of Dr. Chih-Wei Hsu (3)) to minimize dead space and ensure good sealing without excessively tight fitting of the snout. Facemask dead space (about 200 µL) was estimated using the 3D model of the facemask connected to the 3D model of the snout. The pneumotachograph was composed of a unicapillary sensor (inner diameter: 0.9 mm, length: 16 mm, diameter of pressure ports: 0.5 mm, distance between pressure ports: 4.3 mm) connected directly to a miniature pressure transducer (SDP37, Sensirion, Staefa, Switzerland; range: ± 1.25 cm $H_20$). Pressure signals were digitized at 1000 Hz (USB-6210, National Instruments, Houston, TX, USA), recorded and computed with custom software (LabVIEW, National Instruments, Houston, TX, USA). The pneumotachograph and facemask were combined into one single component (“pneumotachometer") and 3D printed (ProtoLabs Ltd, Le Bourget du Lac, France) in a plastic material (MicroFine Green$^{TM}$) (Figures 1A and 1B). The pneumotachometer showed strong linearity over the 0 to 50 mL/min measurement range and a sensitivity of 0.73 x $10^{-3}$ cm $H_2O$.min/mL (Figure E1). Its resistance was 0.25 cm $H_20$.s/mL, a value much lower that airway resistance in one-day old mice, which was estimated 19.66 ± 3.93 cm $H_2O$.s/mL (4). In order to prevent $CO_2$ rebreathing and water accumulation, a bias flow (20 mL/min) was injected into the facemask through a side tube and ran out through the pneumotachometer. The bias flow value was empirically set based on mean ventilation ($V_E$) of newborn mouse pups and the volume of the mask dead space. The dynamic response of the pneumotachometer was analyzed using a sinusoidal flow waveform (amplitude 15 µL and frequency from 0.5 to 10 Hz) provided by a built-in pump incorporating a 50 µL precision syringe (MS*GFN50, Ito Corporation, Tokyo, Japan). The pump was connected to the facemask through a 3D printed model of the snout attached to the facemask. At each frequency, volume estimation error was calculated as (100 x ($V_{meas}$ - $V_{ref}$) / $V_{ref}$), with $V_{ref}$ = 15 µL and $V_{meas}$ as the amplitude of the volume signal calculated by integration of the flow signal. The volume estimation error remained between -2% and 1.5% over the frequency range. The pneumotachometer was calibrated before each experimental session with a 50 µL precision syringe (MS*GFN50, Ito Corporation, Tokyo, Japan) operated manually. The hypercapnic challenge was achieved by switching the air flow through the facemask to an hypercapnic mixture (5 min at 8% $CO_2$, 20.9% $O_2$, 71.1% $N_2$). The washout of gases within the mask after each switch was estimated to take less than five seconds.

**Laser profilometer**

Abdominal movements were detected using a laser profilometer (LJ-V7080 head with a LJ-V7001 controller, Keyence Corporation, Osaka, Japan) pointing radially at a 2 mm x 0.05 mm zone of the lateral abdominal wall (Figures 1A and 1B).

**Electrophysiology**

For immunolabeling of hypoglossal motoneurons we used a mouse anti-islet1,2 (1/250; DSHB) primary antibody and a Goat anti-mouse AlexaFluor 488 (1/500; ThermoFisher) secondary antibody. In electrophysiological recordings from isolated brainstem slice preparations, measurements were taken from 15 consecutive bursts per preparation to determine burst characteristics. Burst rise time was measured from the activity onset to the maximum amplitude (i.e., discharge intensity) of each burst. Burst duration was measured between the onset of the burst and the return to base line. The delay between hypoglossal and phrenic activities was measured from the onset of the phrenic burst.

**SUPPLEMENTAL FIGURE E1**

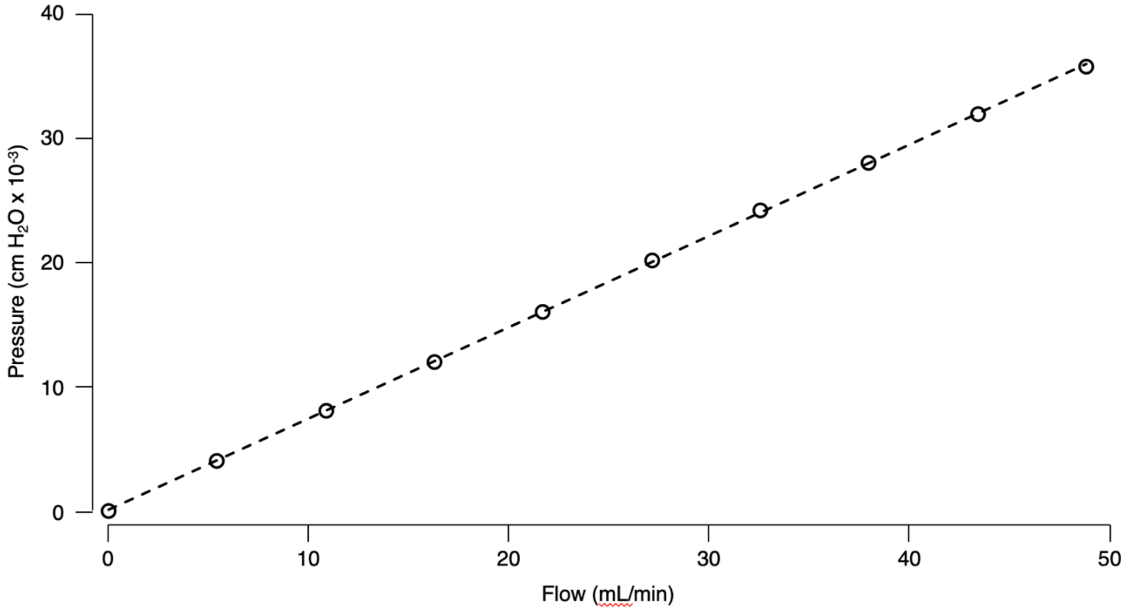


**Figure E1.** Pressure-flow relationship of the pneumotachometer. Stationary flows were successively generated for 10 s by a calibrated mass flow controller (5850S**,** Brooks Instruments, Hatfield PA, USA) connected to the pneumotachometer. Flow values were corrected for temperature, pressure and humidity at experimental conditions. The equation of the least squares regression line was: Pressure = 0.73 x Flow + 0.11. Note the strong linearity of the pressure-flow relationship over the measurement range with $R^2 > 0.999$.

**SUPPLEMENTAL FIGURE E2**

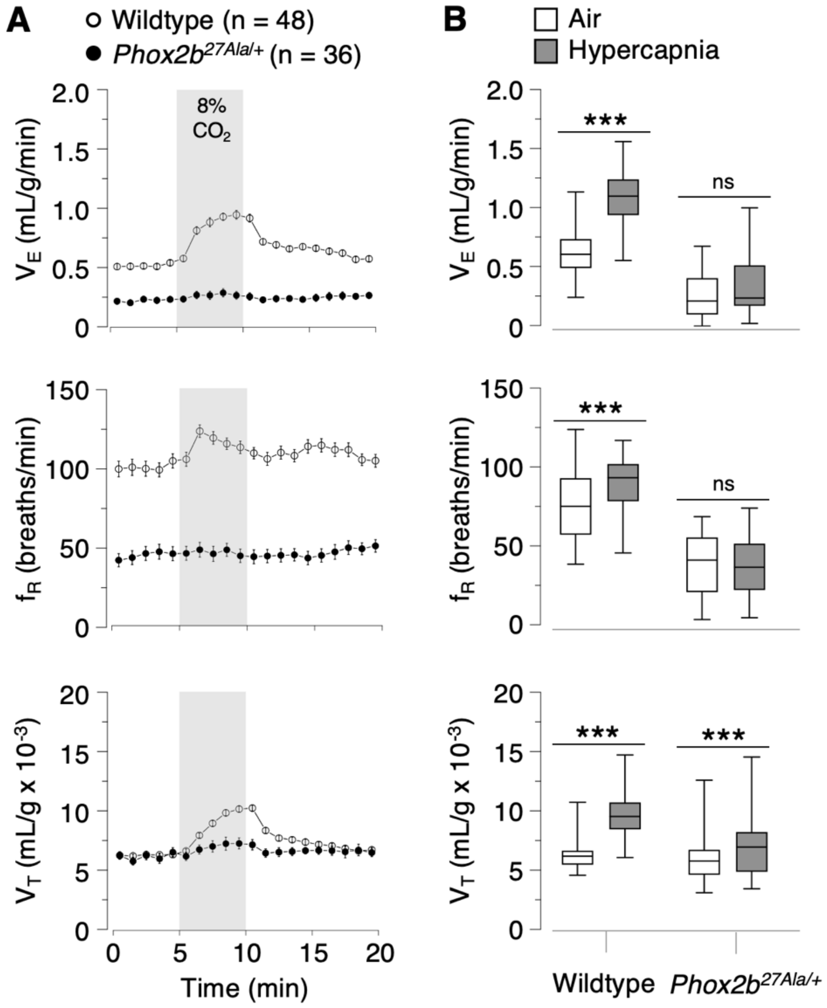


**Figure E2.** Absence of ventilatory response to hypercapnia in *Phox2b*$^{27Ala/+}$ pups (n = 36) unlike in wildtypes (n = 48). (*A*) $V_E$: ventilation; $f_R$: breathing frequency; $V_T$: tidal volume. Values are means ± SEM. (*B*) Lines and whiskers in boxplots represent the medians, interquartile ranges, and minimum and maximum values. Air: average of values measured over 3 min before switching to 8% $CO_2$. Hypercapnia: average of values measured over the last 3-min period of hypercapnia. The small $V_T$-response to $CO_2$ in *Phox2b*$^{27Ala/+}$ pups did not translate into significant $V_E$-increase. In wildtypes, the strong $f_R$ and $V_T$-responses resulted in vigorous $V_E$-increases. Genotype (wildtype vs. *Phox2b*$^{27Ala/+}$) by stimulus (air vs. $CO_2$) interactions: $V_E$: $P < 0.0001$; $f_R$: $P < 0.01$; $V_T$: $P < 0.0001$.

**SUPPLEMENTAL FIGURE E3**

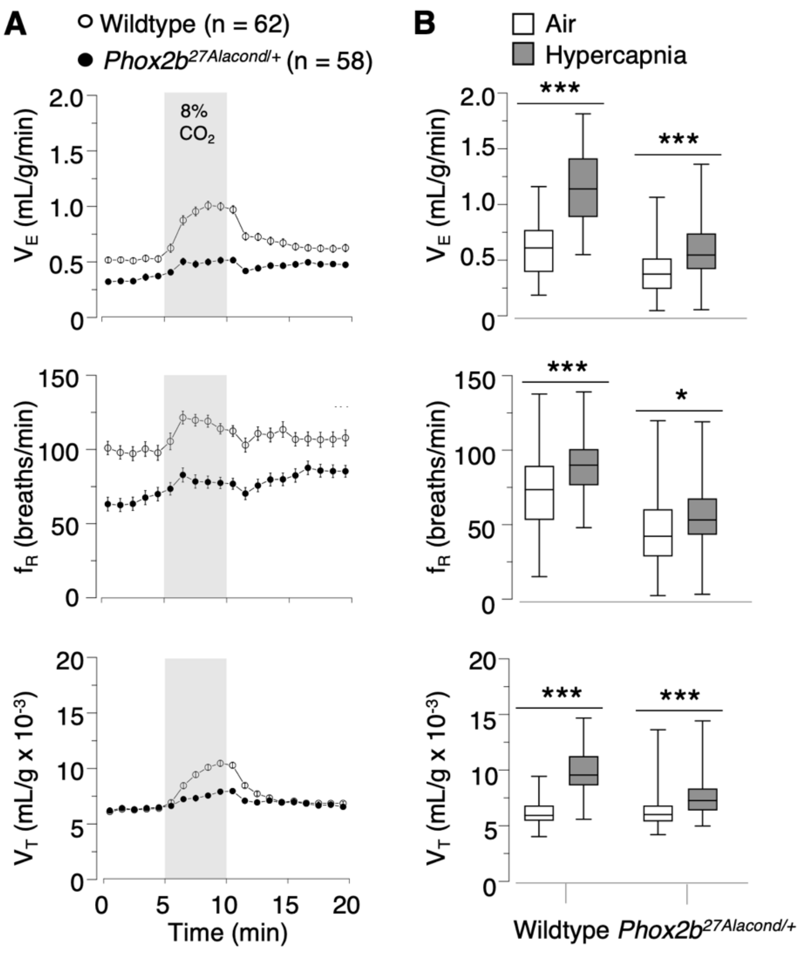


**Figure E3.** Markedly depressed ventilatory response to hypercapnia in *Phox2b*$^{27Alacond/+}$ pups (n = 58) compared to wildtypes (n = 62). (*A*) $V_E$: ventilation; $f_R$: breathing frequency; $V_T$: tidal volume. Values are means ± SEM. (*B*) Lines and whiskers in boxplots represent the medians, interquartile ranges, and minimum and maximum values. Air: average of measured values over 3 min before switching to 8% $CO_2$. Hypercapnia: average of measured values over the last 3-min of hypercapnia. Both groups significantly increased $V_E$ in response to 8% $CO_2$, but the increase was significantly larger in wildtypes. Genotype (wildtype vs. *Phox2b*$^{27Alacond/+}$) by stimulus (air vs. $CO_2$) interactions: $V_E$: $P < 0.0001$; $f_R$: $P < 0.1$, not significant; $V_T$: $P < 0.0001$. At variance with a previous study in which the $V_E$-response to hypercapnia was reported to be fully abolished in mutants, the small response found here may reflect a better accuracy of pneumotachography compared to whole-body plethysmography.
